## Supplementary for "mcRigor: a statistical method to enhance the rigor of metacell partitioning in single-cell data analysis"

### Supplementary Figures

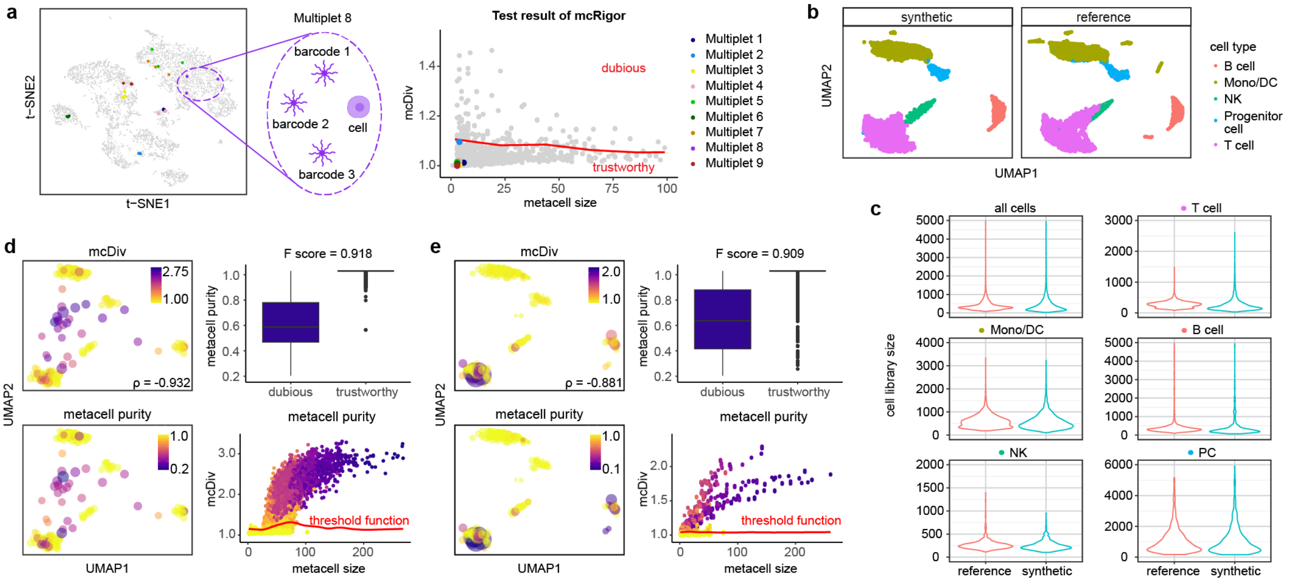


**Supplementary Fig 1: Justification of the mcRigor method.** **a,** Barcode multiplets justify the statistical definition of metacell (left) and the reliability of mcRigor (right). **b,** UMAP plots of the semi-synthetic data and the reference real data. **c,** Violin plots of cell library sizes for the semi-synthetic data versus the reference data. **d,** mcRigor effectively measures metacell heterogeneity and detects dubious metacells from the SEACells partition on the semi-synthetic data. Left: UMAP plots of the metacells colored by mcDiv values versus metacell purity (with Pearson correlation $\rho = -0.932$). Top right: mcRigor distinguishes between ground-truth dubious and trustworthy metacells with high accuracy (F-score = 0.918). Bottom left: The scatter plot of mcDiv versus metacell size with the dubious metacell detection threshold function. **e,** mcRigor effectively measures metacell heterogeneity and detects dubious metacells from the SuperCell partition on the semi-synthetic data. Similar to **d**, the Pearson correlation is $\rho = -0.881$, and the F-score is 0.909.

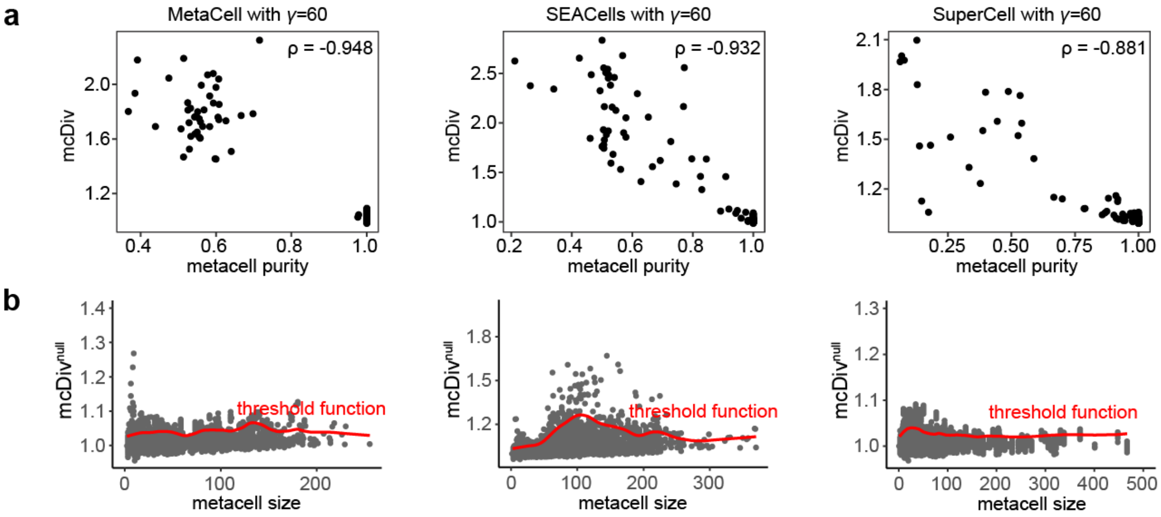


**Supplementary Fig 2: Additional simulation results of mcRigor on the semi-synthetic dataset. a,** The mcDiv statistic is highly correlated with true metacell purity. $\rho$ stands for Pearson correlation. **b,** Scatter plots showing the distributions of the null statistic value, $\text{mcDiv}^{\text{null}}$, contructed by mcRigor, and the metacell-size-specific threshold function for detecting dubious metacells (as those below their size-specific thresholds).

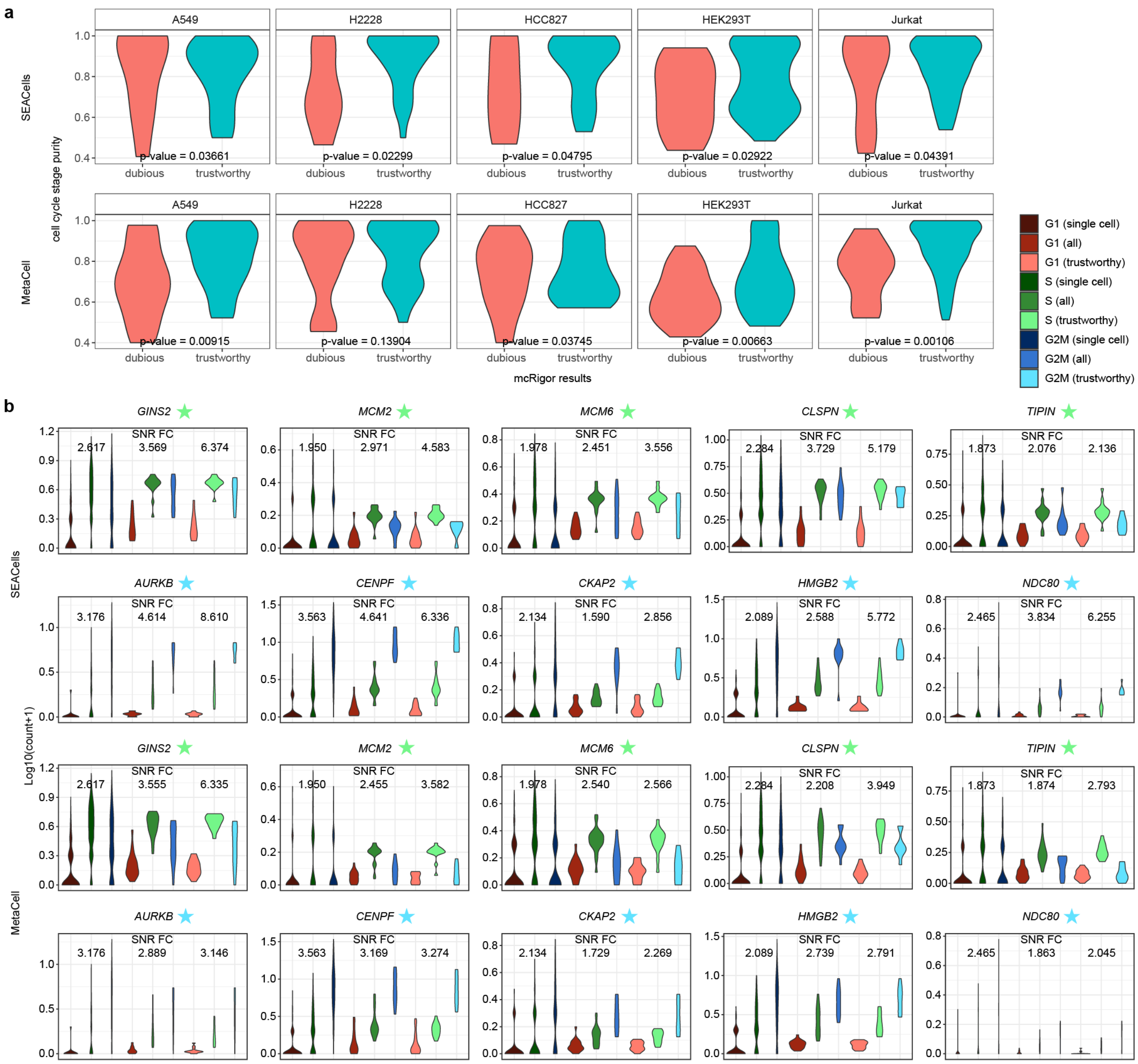


**Supplementary Fig 3: mcRigor's trustworthy metacells reveal cell-cycle phases within cell lines.** **a,** Violin plots comparing the cell cycle-phase purity distributions of dubious metacells and trustworthy metacells. Trustworthy metacells consistently exhibit higher purity than dubious metacells. **b,** Violin plots displaying the $\log_{10}$(count+1) expression levels of 10 cell-cycle marker genes across single cells, all metacells ("all"), and trustworthy metacells ("trustworthy"). The metacells were generated by two metacell methods: SEACells and MetaCell, with $\gamma = 20$. SNR FC represents the fold change in signal-to-noise ratio for the phase associated with each marker gene (indicated by a star) relative to the other two phases.

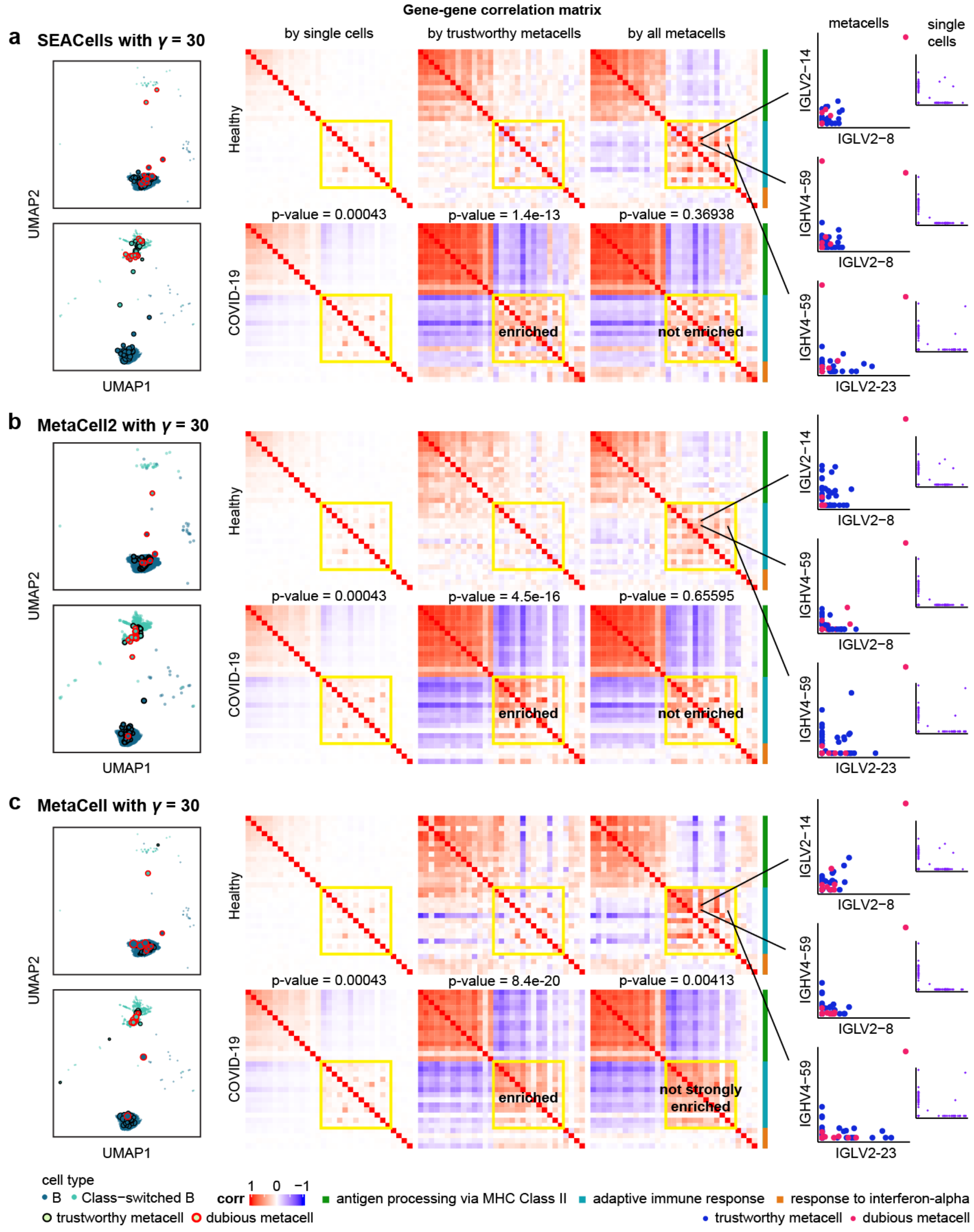


**Supplementary Fig 4: mcRigor reveals gene co-expression modules enriched for COVID-19 compared to healthy control by removing signal distortion caused by dubious metacells.** This is a continuation of Fig 1e in the main text with analysis results for three more metacell methods: SEACells (**a**), MetaCell2 (**b**), and MetaCell (**c**).

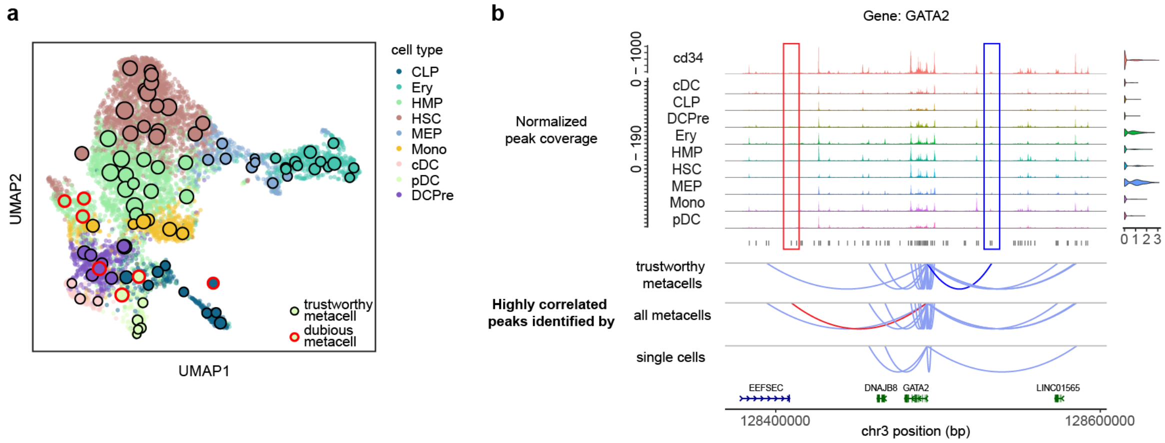


**Supplementary Fig 5: mcRigor identifies dubious metacells over the original SEACells partitioning and rectifies gene regulatory inference by removing the dubious metacells. a,** UMAP plot showing dubious metacells detected by mcRigor from the original SEACells partitioning. **b,** Highly correlated peaks for gene *GATA2* identified using trustworthy metacells, all metacells, or single cells. This is the complete version for the right panel of Fig 1f (right) in the main text, where we only showed peak signal coverage plots for three cell types (Ery, HSC, and MEP).

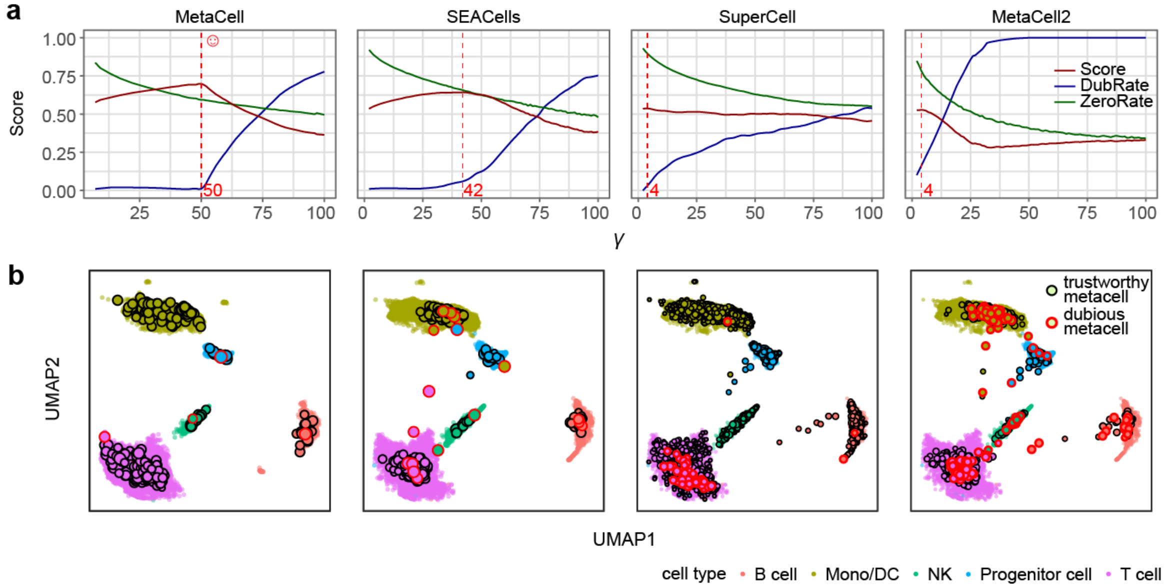


**Supplementary Fig 6: mcRigor optimizes metacell partitioning for the semi-synthetic dataset. a,** Line plots showing the evaluation scores provided by mcRigor. The vertical red dashed lines mark the optimal $\gamma$ selected for each metacell method. The red smiling face marks the optimal metacell partition selected across all method-hyperparameter configurations. **b,** Single-cell UMAP plots showing the optimal metacell partitioning for each method, with dubious metacells highlighted in red circles.

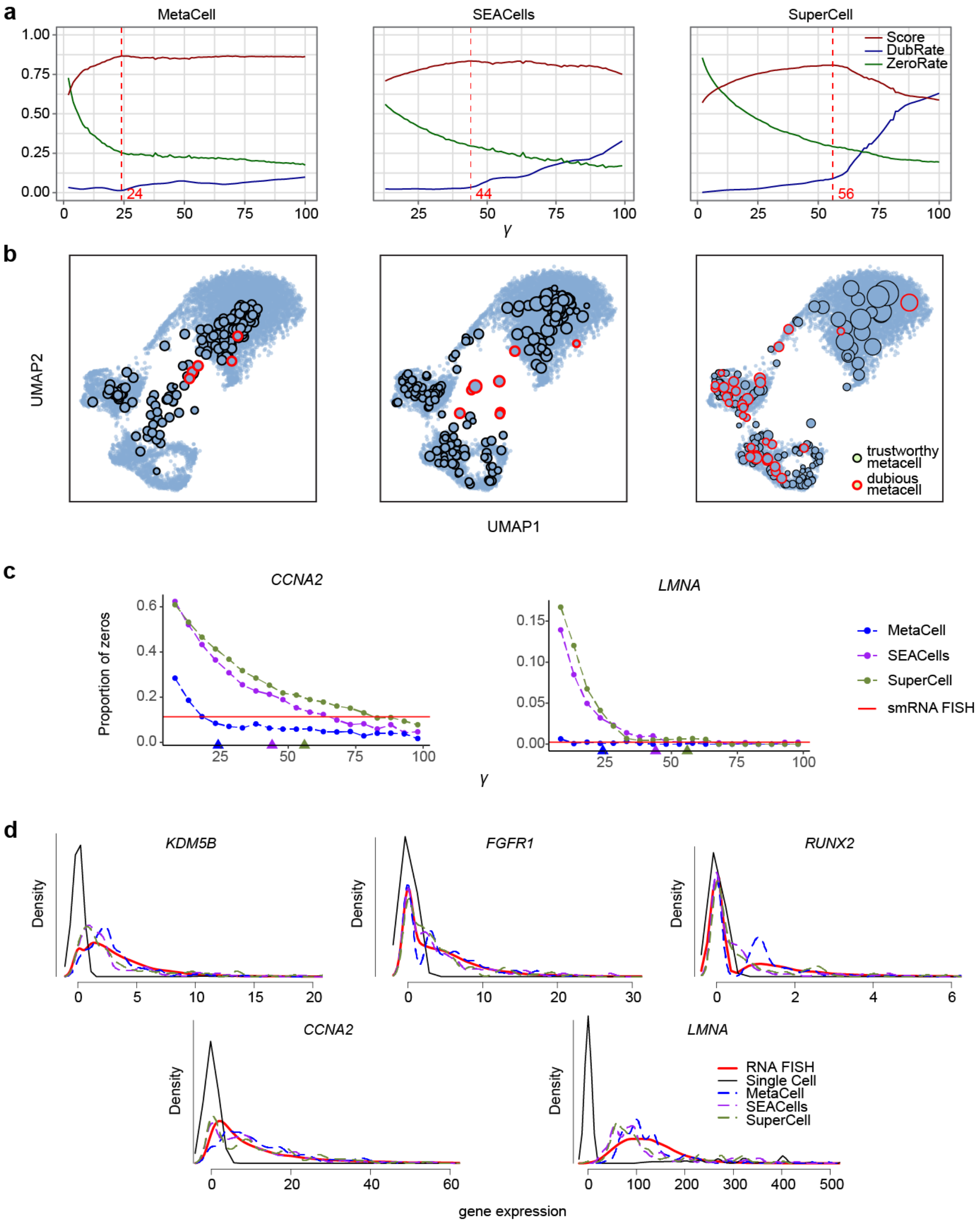


**Supplementary Fig 7: Additonal results from applying mcRigor to the Drop-seq + smRNA FISH dataset. a,** The evaluation scores provided by mcRigor. The vertical red dashed lines mark the optimal $\gamma$ selected for each metacell method. **b,** Single-cell UMAP plots showing the optimal metacell partitioning for each method, with dubious metacells highlighted in red circles. **c,** Line plots showing zero proportions for genes *CCNA2* and *LMNA* under metacell partitionings generated by the three metacell methods with varying $\gamma$ values (with triangles indicating the optimal $\gamma$ values selected in **a**). The red horizontal line marks the zero proportion in the smRNA FISH data. **d,** Density plots showing gene expression from the single cell profiles, metacell profiles (generated by the optimal $\gamma$ for each metacell method), and the smRNA FISH profiles.

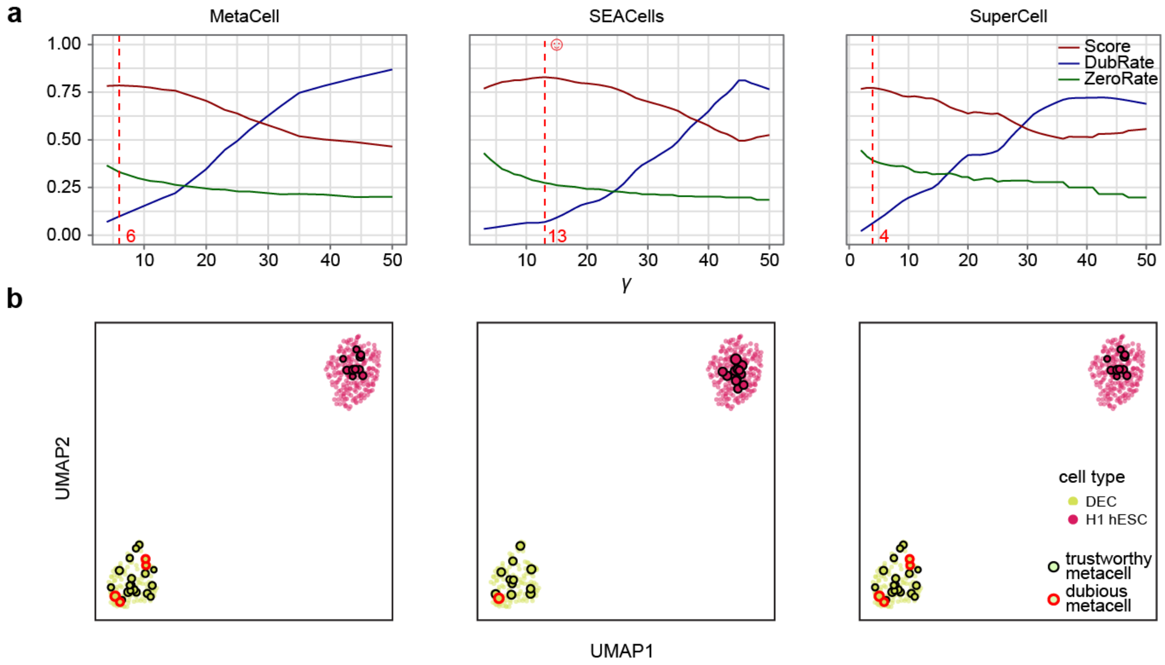


**Supplementary Fig 8: Additonal results from applying mcRigor to the scRNA-seq + bulk ESC dataset. a,** The evaluation scores provided by mcRigor. The vertical red dashed lines mark the optimal $\gamma$ selected for each metacell method. The red smiling face marks the optimal metacell partition selected across all method-hyperparameter configurations. **b,** Single-cell UMAP plots showing the optimal metacell partitioning for each method, with dubious metacells highlighted in red circles.

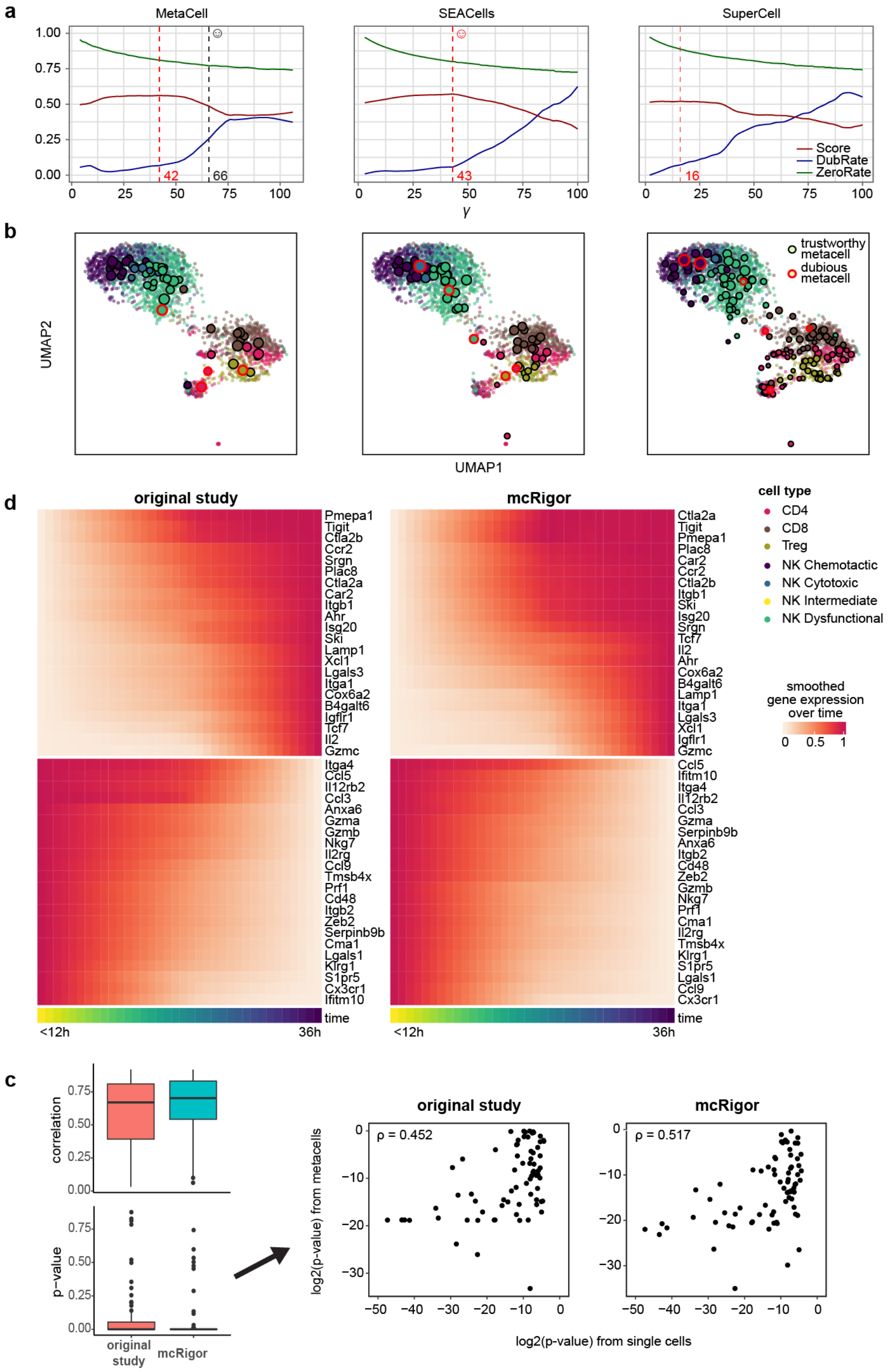


**Supplementary Fig 9: Additonal results from applying mcRigor to the Zman-seq dataset.** **a,** The evaluation scores provided by mcRigor. The vertical red dashed lines mark the optimal $\gamma$ selected for each metacell method. The red smiling face marks the optimal metacell partition selected by mcRigor across all method-hyperparameter configurations. The black smiling face marks the metacell partition used in the original study. **b,** Single-cell UMAP plots showing the optimal metacell partitioning for each method, with dubious metacells highlighted in red circles. **c (bottom row),** The single-cell DE genes exhibited higher correlations with tumor exposure time and lower p-values when using the optimal metacell partition selected by mcRigor compared to the original partition. **d (third row),** The optimal metacell partition selected by mcRigor recovers the time-dependent genes identified in the original study.

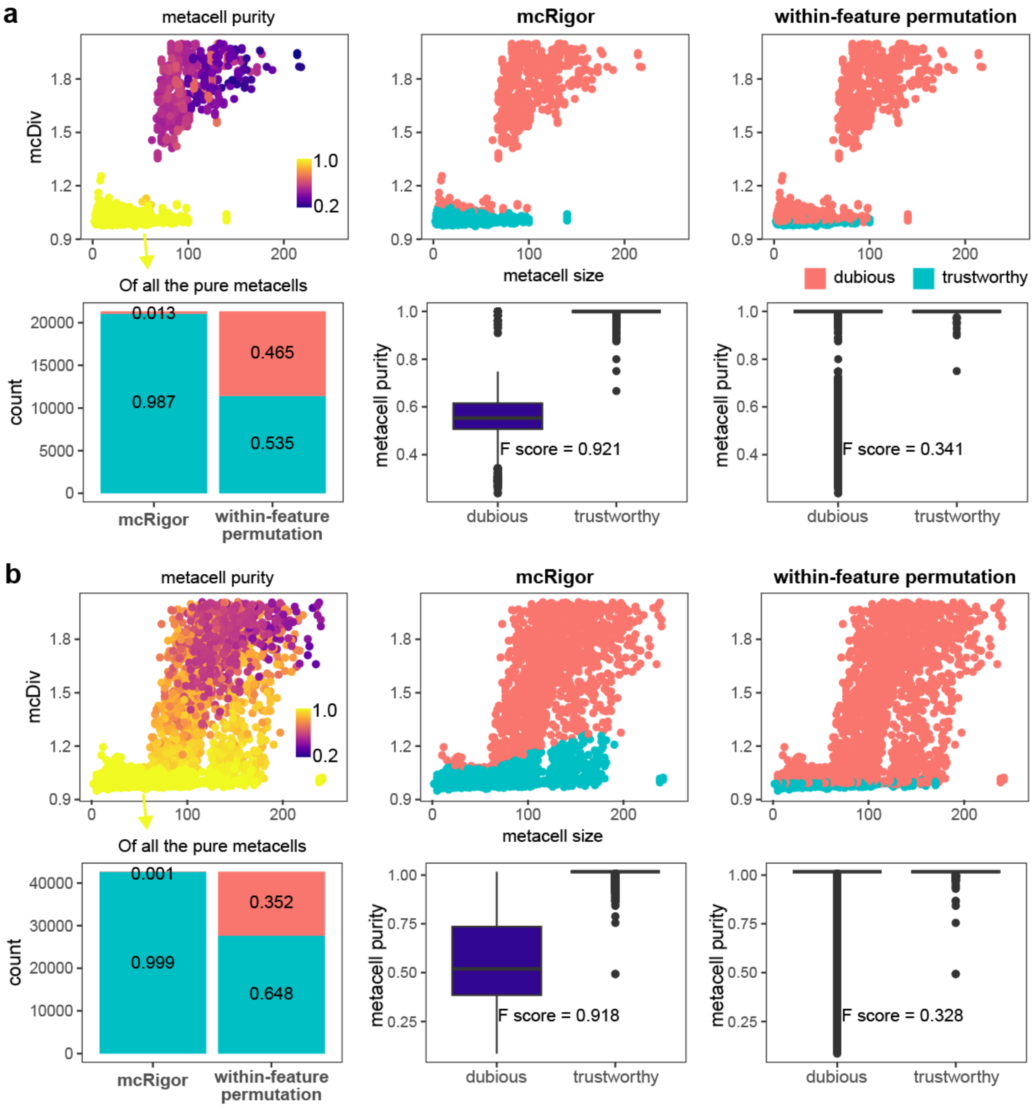


**Supplementary Fig 10: Within-feature permutation alone is not a valid null for dubious metacell detection.** **a,** Comparison of dubious metacell detection results obtained by applying mcRigor (double permutation) and within-feaure permutation to the metacell partition generated by MetaCell on the semi-synthetic data. **b,** Comparison of dubious metacell detection results obtained by applying mcRigor (double permutation) and within-feaure permuation to the metacell partition generated by SEACells on the semi-synthetic data.

### Distribution of metacell sizes

We observed that, at the same granularity level, metacells display substantial variability in size (Fig. 1c, Supplementary Fig. 5a, Supplementary Fig. 9b). To further investigate how metacell sizes are distributed, we plotted histograms of metacell sizes for the various cell types in the datasets we analyzed (Supplementary Fig 11, Supplementary Fig 12). Interestingly, the distributions of metacell sizes differ considerably across cell types. This variation may arise from certain biological states being more stable, thereby encompassing more cells and forming larger metacells, while less stable states include fewer cells, leading to smaller metacells. For instance, in the `bmcite` dataset, progenitor cells are generally grouped into smaller metacells, whereas T cells are aggregated into larger metacells (Supplementary Fig 12a). Furthermore, metacell size distributions vary by condition: in the COVID-19 PBMC dataset, small metacells are more abundant in the COVID-19 group than in the healthy group (Supplementary Fig 11a), indicating less stable biological states under the diseased condition.

We also observed no clear relationship between metacell size and trustworthiness (as determined by mcRigor), indicating that dubious metacells cannot be reliably identified based solely on size.

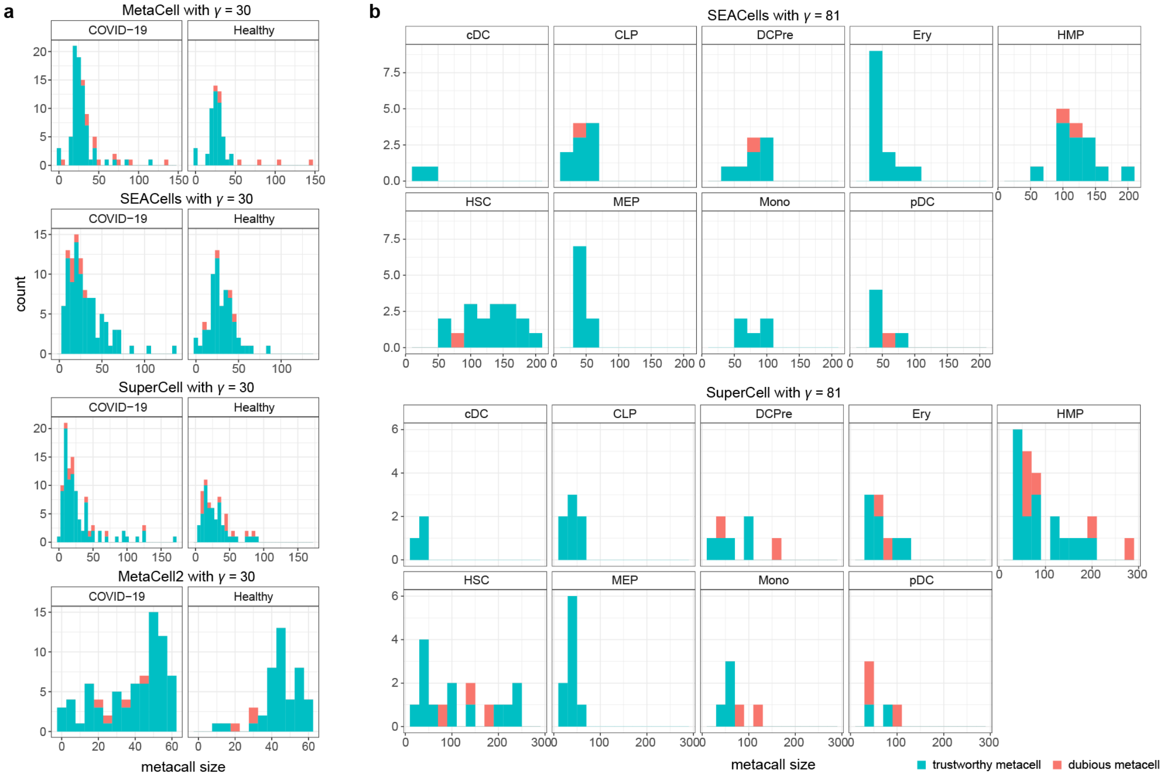


**Supplementary Fig 11: Distribution of metacell sizes** for (**a**) different conditions (COVID-19 versus Healthy) in the COVID-19 PBMC dataset, and (**b**) different cell types in the scMultiome dataset.

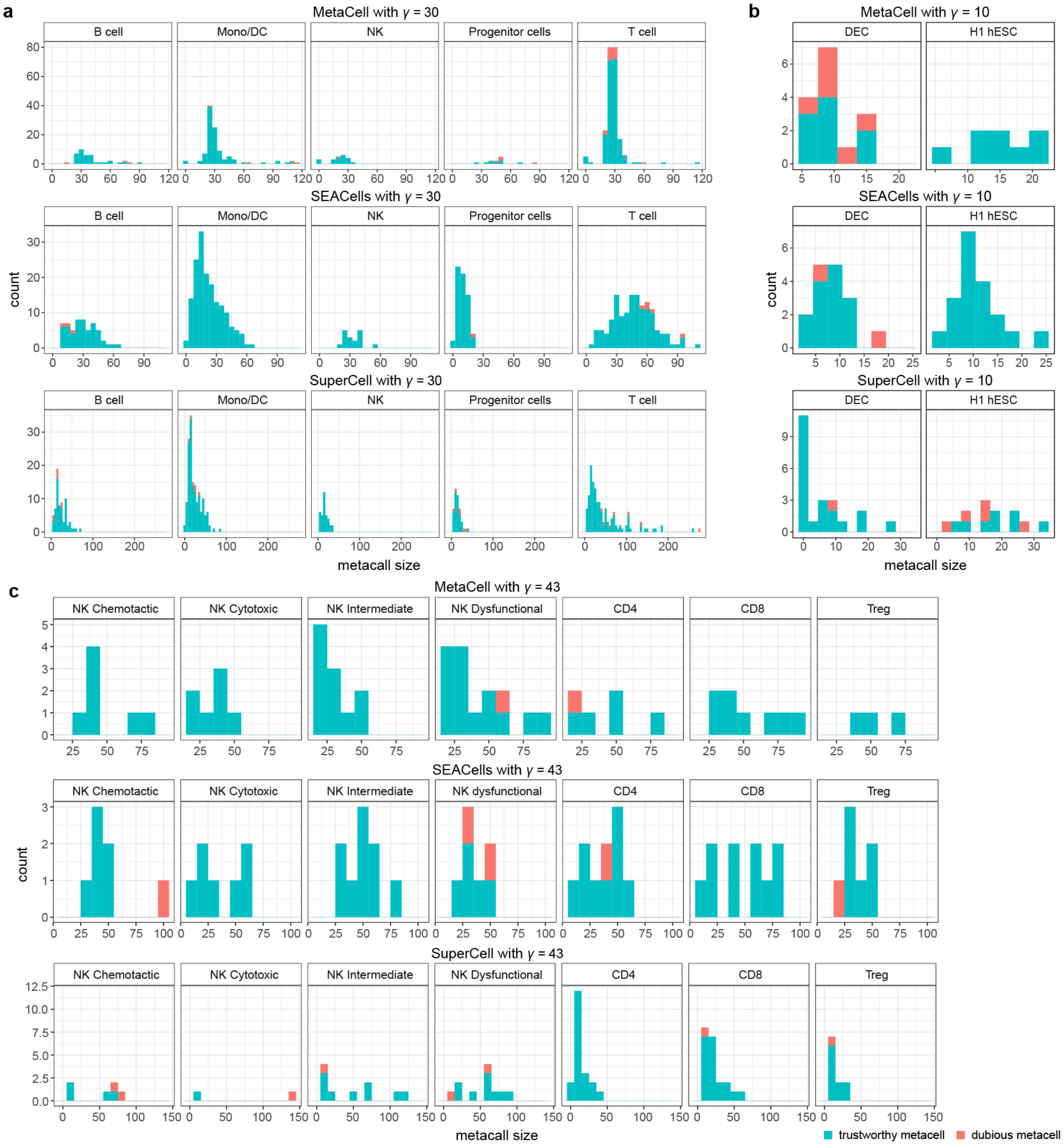


**Supplementary Fig 12: Distribution of metacell sizes for different cell types** in (**a**) the `bmcite` dataset, (**b**) the scRNA-seq + bulk ESC dataset, and (**c**) the Zman-seq dataset.
